# Genotype-dependent and heat-induced grain chalkiness in rice correlates with the expression patterns of starch biosynthesis genes

**DOI:** 10.1101/2020.10.16.342873

**Authors:** Peter James Gann, Manuel Esguerra, Paul Allen Counce, Vibha Srivastava

**Affiliations:** Cell and Molecular Biology Program, University of Arkansas, Fayetteville; Department of Crop, Soil and Environmental Sciences, University of Arkansas, Fayetteville; Rice Research and Extension Center, Stuttgart, Arkansas; Department of Horticulture, University of Arkansas, Fayetteville

**Keywords:** Chalkiness, Starch Biosynthesis, Rice Grain, High Nighttime Temperature

## Abstract

To understand the molecular basis of environment-induced and genotype-dependent chalkiness, six rice genotypes showing variable chalk levels were subjected to gene expression analysis during reproductive stages. In the high chalk genotypes, the peak expressions of *ADP-Glucose Pyrophosphorylase (AGPase) Large Subunit 4* (*AGPL4*) occurred in the stages before grain filling commenced, creating a temporal gap with the upregulation of *Granule Bound Starch Synthase I* (*GBSSI*) and *Starch Synthase IIA* (*SSIIA*). Whereas, in the low chalk genotypes, *AGPL4* expression generally occurred in later stages, close to the upregulation of *GBSSI* and *SSIIA*. However, heat treatment altered the expression pattern and created a gap between the expression peaks of *AGPL4*, and *GBSS1* and *SSIIA*. This change was accompanied by transformed granular morphology, increased protein content, and chalkiness in the grains. *AGPL4* expression pattern may partially explain chalkiness as it contributes to the pool of ADP-Glucose for producing amylose and amylopectin, the major components of the starch. Down-regulation of AGPase during grain filling stages could result in a limited pool of ADP-Glucose leading to inefficient grain filling and air pockets that contribute to chalkiness. The study suggests a mechanism of grain chalkiness based on the coordination of the three starch biosynthesis genes in rice.

**Significance statement:** Genotype-dependent and heat-induced grain chalkiness in rice is partially based on the increased gap between the upregulation *AGPase* and that of *GBSSI* and *SSIIA* through reproductive stages. This temporal gap could limit starch accumulation and alter granular morphology, eventually leading to grain chalkiness.

## INTRODUCTION

Starch is the primary carbon and energy reserve in rice endosperm. Physical, granular and chemical properties are the measures of grain quality, which are dependent on the starch biosynthesis process. Grain chalkiness is a highly undesirable trait in rice, which is genotype-dependent and induced by the high nighttime temperature (HNT) (Feng et al. 2017; Jagdish et al. 2015; Lanning et al. 2011; Xu et al. 2020). Coordination of the enzymes involved in this process is important to prevent chalk formation in the grain that affects the market value, cooking and eating quality of rice (Fitzgerald & Resurreccion 2009; Lisle et al. 2000).

Briefly, starch biosynthesis starts after fertilization, when the endosperm cells multiply, form cell walls and elongate. Prior to starch biosynthesis, cell walls must be present along with the appropriate contingent of cell organelles. Formation and elongation of cell walls utilizes imported sucrose. There are three sucrolytic enzymes that breakdown sucrose in the endosperm namely acid invertase, neutral invertase and sucrose synthase. However, early cell wall development is largely dependent on the action of acid invertase to convert the sucrose into glucose and fructose. Consequently, sucrose is also catalyzed by neutral Cell Wall Invertases (*OsCIN*) (Abu-Zaitoon et al. 2012; Wang et al. 2008). Besides contributing to cell wall development, vacuolar acid invertase (*INV3*), sucrose synthase (*RSUS1*) also contributes to the elongation of the cell (Hirose, et al. 2004; Ishimaru et al. 2005). After the cell wall and organelles form, the grain begins to fill.

During grain filling, UDP-glucose, glucose and fructose, metabolic products of sucrolytic enzymes, are converted to Glucose-1-Phosphate by several enzymes. ADP-glucose pyrophosphorylase (AGPase) catalyzes the production of ADP-glucose from Glucose-1-Phosphate. Subsequently, two separate biochemical pathways for adding glucose to α-glucan chains of amylopectin by starch synthases (SS) and to amylose via Granule Bound Starch Synthases (GBSS) commence (Pfister & Zeeman, 2016). AGPase consists of four subunits, *AGPL2* and *AGPL4* as the regulatory large subunits, and *AGPS1* and *AGPS2* as the catalytic small subunits (Tuncel et al. 2014). AGPase catalyzes the rate-limiting reaction converting Glucose-1-Phosphate and ATP to ADP-Glucose and Pyrophosphate (Iglesias & Preiss,1992; Sivak & Preiss, 1998). With the reversibility of the AGPase-catalyzed reaction, the follow up expression of GBSS and SS is important to utilize ADP-Glucose and prevent the futile cycle of converting ADP-Glucose back to Glucose-1-Phosphate (Barratt et al. 2009 and Baroja-Fernandez et al. 2012).

ADP-Glucose serves as the substrate for GBSS and SS, two enzymes with multiple isoforms. GBSS synthesizes amylose with α(1 **→** 4) glycoside linkage, and SS elongates polysaccharide chain with α (1 **→** 4 & 1 **→** 6) glycoside linkages synthesizing amylopectin. *GBSSI and SSIIA* are the highly expressed isoforms in the rice endosperm during grain filling stages (Hirose et al. 2004; Ohdan et al. 2005; Umemoto et al. 2002; Xing et al. 2016). Mutations in starch synthase genes have been described to affect grain chalkiness by altering the granular morphology from compound polyhedral type to simple spherical type (Toyosawa et al. 2016; Kusano et al. 2012). The simple spherical granules constitute the chalk portion as they pack loosely and include airspaces (Kaneko et al. 2016; Kim et al. 2004; Lu et al. 2015; Mitsui et al. 2016). Thus, starch synthesis genes play a major role in determining grain quality; however, their coordination with one another and expression patterns related with chalky or translucent grains has not been fully understood.

Some of the other mechanisms that control grain chalkiness are related to the starch accumulation process. For example, disruption of the amyloplast’s outer envelope membrane (OEM) during seed maturation leads to the formation of simple and spherical granules (Toyosawa et al. 2016), and early degradation of starch through amylase activity contributes to grain chalkiness. Micropores on the surfaces of rough amyloplast in the chalky grains indicate starch degradation by amylase activities (Lin et al. 2016). Finally, protein bodies in the endosperm are also correlated with chalkiness. Several studies showed that chalky rice contains abnormal protein bodies in the endosperm that are large in size and accommodate more air spaces (Nagamine et al. 2011; Ren et al. 2014; and Fukuda et al. 2011).

In this study, rice genotypes consisting of well-known chalky varieties and low chalk cultivars were subjected to gene expression analysis as well as the analysis of grain physical characteristics, starch components, and the granular morphology. The expression pattern of *AGPL4*, *GBSSI* and *SSIIA* was found to be genotype-dependent and heat-sensitive, which highlights the importance of coordinated starch biosynthesis during the critical stages of grain filling to produce properly filled non-chalky, translucent rice grains. The disruption in the expression pattern of starch biosynthesis genes by heat appears to be a part of the mechanism associated with the environment-induced chalkiness.

## MATERIALS AND METHODS

### Plant materials

Six genotypes, ZHE 733, Nagina 22 (N22), Nipponbare (Nip), Taggart, Diamond, and LaGrue, representing *indica*, *aus* or *japonica* sub-species were used in this study. These genotypes included 3 cultivars (Taggart, Diamond, and LaGrue) developed at Arkansas Rice Research Center. Three replications of each genotype were planted on July 2019 in the greenhouse. When plants were at R0 or R1 stage (Moldenhauer et al. 2018), they were transferred to growth chambers. For the normal condition, the temperature was set at 30°C day / 22°C night, and for heat-stress at 30°C day / 28°C night with nighttime starting at 8PM and ending at 6AM. Relative humidity and lighting conditions were uniform for the two set-ups. The rice plant culms entering the reproductive stage were tagged, and used as the source of samples for the different stages, namely before panicle emergence (BP) (R2), early flowering/after panicle emergence (AP) (R3 or R4), 5 days after flowering (DAF) (R5 to R6), 10DAF(R6), 15DAF (R8) and 20DAF (R8) (Moldenhauer et. al., 2018). For the granular, physical, and chemical properties, grains from the second panicle were collected at 25DAF. Immediately after collection, the samples for gene expression analysis were frozen in liquid nitrogen and stored at –8°C.

### Grain physical property

Grains collected at 25DAF were dried under room temperature for two weeks after harvesting, prior to the observation for chalkiness, and powdered by a cyclone milling machine for the chemical property analysis. Chalkiness was measured using WinSEEDLETM with 150 grains for each genotype. The percentage of chalky grains and the average chalk size per grain were taken as measurements of chalkiness.

### Gene expression from databases

Heatmaps were generated to select the appropriate genes for the starch biosynthetic pathway of interest. RiceExpro was used under the category datasets and gene expression profile at different ripening stages (7DAF, 10DAF, 14DAF, 21DAF, 28DAF, and 42DAF) relevant to the stages selected for gene expression analysis by qPCR.

### Transcript levels

Total RNA was isolated using Trizol (Invitrogen Inc.) and quantified using Nano-drop 2000 (Thermo-Fisher Inc). Two micrograms of total RNA were treated with RQ1-RNAse free DNase (Thermofisher Inc.), and one microgram of the DNase-treated RNA was used for cDNA synthesis using PrimeScript RT reagent kit (Takara Bio, CA, USA). The expression analysis was performed using TB green Premix Ex Taq II (Takara Bio, CA, USA) on Bio-Rad CFX 96 C1000 with following conditions: 95°C for 30 sec. and 40 cycles of 95°C for 5 sec + 60°C for 30 sec. The product specificity was verified by the melt curve analysis. The Ct values of genes-of-interest were normalized against *7Ubiquitin fused protein* as the reference gene.

### Granular morphology

Grains were split in two using a microtome for the cross-section perspective. Cross-section of the grains were viewed under Philips/Fei XL-30 Environmental Scanning Electron Microscope (ESEM) with the settings Acc V. of 10kV, 2000x magnification, 3.0 spot and 10 μm bar. The surfaces of whole grains were captured both in low and high magnifications. Captured images were adjusted to brightness of 20, color balance of R (0), G (0) and B (−20), and gamma correction of 1.00.

### Biochemical Analysis

The amylose and amylopectin content of grains were determined using the Megazyme amylose/amylopectin assay (K-AMYL) following the manufacturer’s method. Protein quantification was done using 50 mg of grain powder in 1 ml of TE buffer pH 8.0 using Bradford assay against a standard curve of BSA (0, 2.5, 5, 7.5, and 10 µg/ml). Absorbance was read at 595 nm in Bio-Rad SmartSpec 3000 spectrophotometer.

### Data analysis

The experiment was conducted in a completely randomized design with three independent replications. Data were subjected to one-way analysis of variance (ANOVA). To determine the significant differences in the amylose and amylopectin content and protein concentration, Tukey’s range test was used to compare the genotypes under normal condition, and Student’s T-test for pairwise comparison in the normal and heat conditions. All statistical analyses were performed in SAS statistical software (version 9.4, SAS Institute Inc., Cary, NC).

## RESULTS AND DISCUSSION

### Expression patterns of starch biosynthesis genes

The heatmap based on the publicly available databases showed genes encoding the four subunits of plastidial *AGPase* are expressed differentially. *AGPS1* and *AGPL2* are consistently expressed, indicating the availability of these subunits throughout the reproductive stages, while *AGPS2* is consistently downregulated. *AGPL4*, on the other hand, is upregulated at 7DAF followed by continual decline in subsequent stages (Figure 1). This pattern suggests that temporal expression of *AGPL4* is critical for plastidial AGPase complex formation. *AGPL1* is a unique subunit in the cytosolic AGPase, which is upregulated until 21DAF (Figure 1). The resulting cytosolic ADP-Glucose passes through the transporters to enter the amyloplast for starch biosynthesis (Pfister & Zeeman, 2016). However, suppression of cytosolic AGPase in barley did not change the granular morphology, an important indicator of chalkiness in rice (Johnson et al. 2003). This suggests that the plastidial AGPase, possibly owing to its *in situ* expression, could have a greater effect on granular morphology and grain chalkiness.

**Figure 1:**
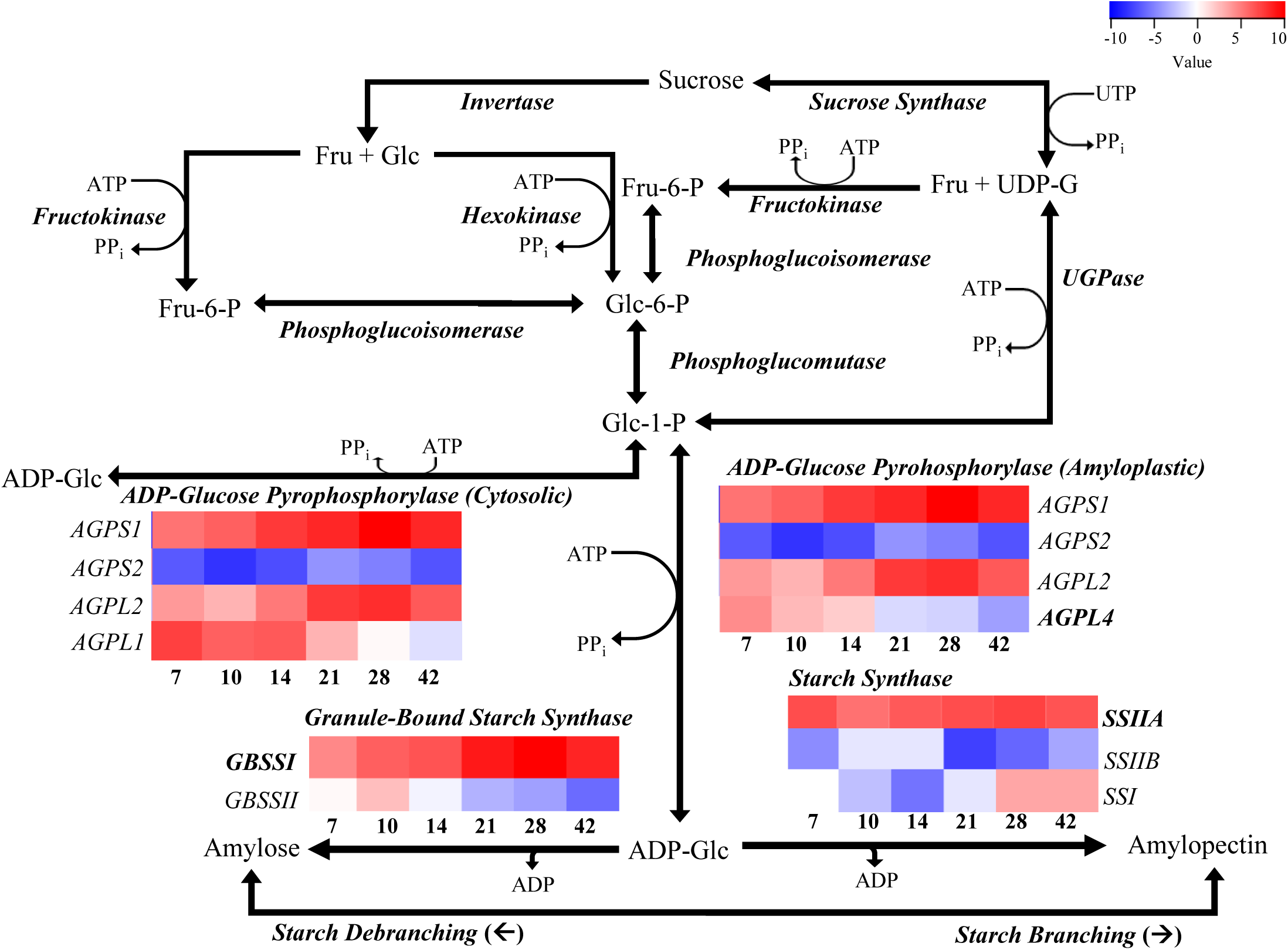
Illustration of the starch biosynthesis pathway. Flow of substrates and products are indicated by the arrows. Genes coding for the enzyme catalyzing a reaction are in bold and italics. Heatmaps for three genes in the two ultimate pathways are placed under the gene name with the specific isoform or subunit indicated. Columns in the heatmap indicate the stages 7 - 42DAF.

Similarly, heatmap of GBSS showed that *GBSSII* is downregulated during advancing stages of grain filling (14DAF onwards), while *GBSSI* is consistently expressed. Further, *SSI* and *SSIIB* are downregulated between 7 – 21DAF, and *SSIIA* is consistently expressed up to 42DAF (Figure 1). These observations matched the previous study, which described *SSIIA* as the steady expresser in all grain filling stages, and *GBSSI* as highly expressed in the mid until the late grain filling stages (Hirose & Terao 2004). Previous studies have also described the roles of differential expression of individual genes in starch biosynthesis. Upregulation of *SSI* at 28 and 42DAF indicates that it plays a role in synthesizing amylopectin during late stages. Corroborating with this, suppression of *SSI* by RNAi decreased amylopectin and altered the granular structure from compound to a simple type in Nipponbare (Zhao et al. 2019). *GBSSI* and *SSIIA* are expressed at higher levels than *AGPL4* throughout the reproductive stages in the endosperm, suggesting a rate limiting effect of *AGPL4*. Several studies have shown that mutations and RNAi suppression of *GBSSI* changes the amylose content in rice (Dobo et al. 2010; Liu et al. 2014; Miura et al. 2018). Variations and deficiency in *SSIIA* have been found to alter amylopectin and starch quality (Nakamura et al. 2005; and Miura et al. 2018). Therefore, *GBSSI* and *SSIIA* play significant roles in starch biosynthesis by controlling amylose and amylopectin content and the granular structure, and a rate limiting effect is possibly imposed by *AGPL4* expression pattern.

### Grain chalkiness in different genotypes

The opaque white in the grains indicates the chalky portion. The six genotypes used in this study were found to have different levels of chalkiness based on which they were classified as high-chalky or low-chalky. High-chalky lines, ZHE 733, Nipponbare, and Nagina 22, contain large opaque areas (Figure 2A-C), while the three low-chalky cultivars, Diamond, Taggart, LaGrue, contain no chalk or small chalky areas (Figure 2D-F). These groups are also distinguished by the frequency at which large or small chalk occur in the grains. In the high-chalky lines, chalk was observed in all grains, with the majority (average of 82%) showing large chalk (>20% of grain size), in addition to small (<10% of grain size) and medium (11- 20% of grain size) size chalk (Figure 3A-C). On the other hand, in low-chalky cultivars, small chalk was found in the majority of the grains (average of 84%) with a small percentage (2-4%) showing no chalk (Figure 3D-F). Among the low-chalky cultivars, LaGrue (Figure 3F) was found to contain more chalk (medium sizes) than Taggart or Diamond (Figure 3D-E).

**Figure 2:**
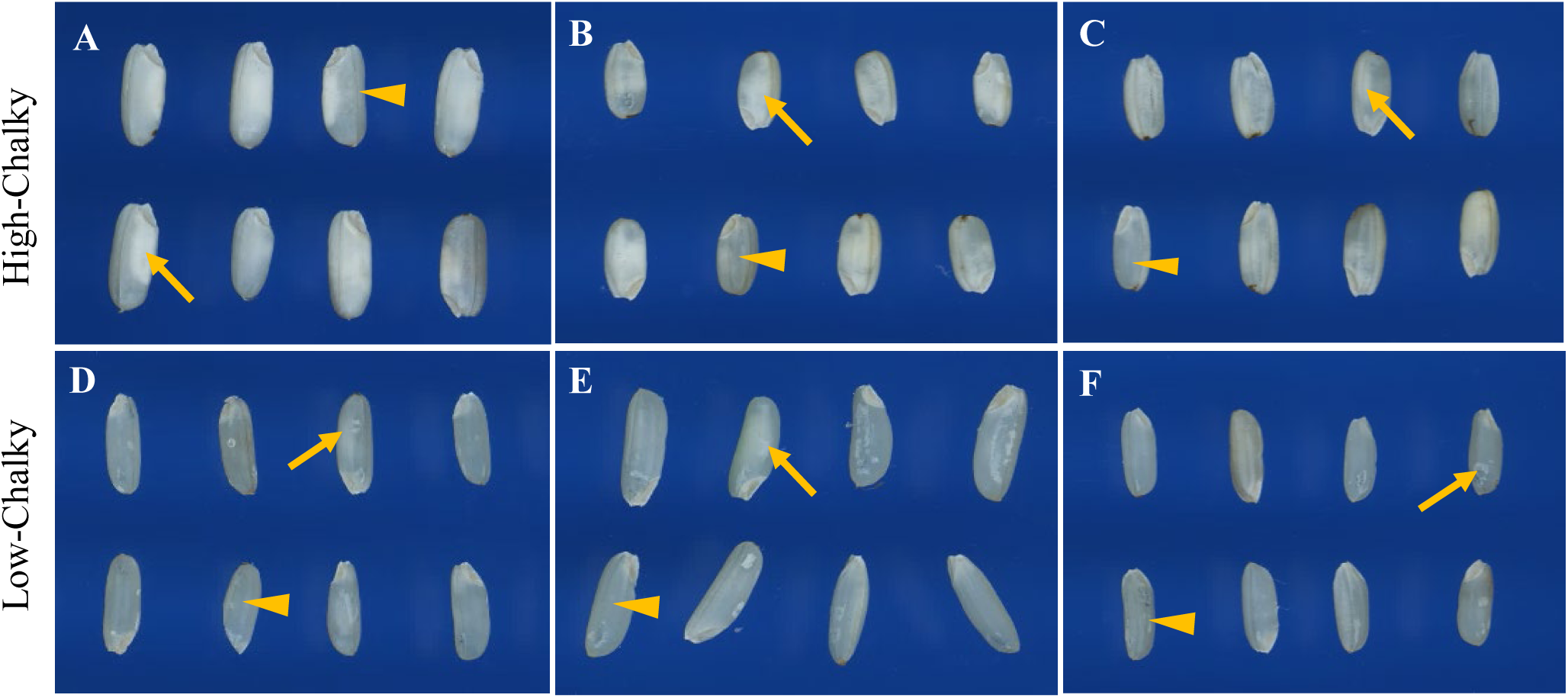
Physical characteristics of rice grains of different genotypes using WinSEEDLE. Arrow points to the chalky area and arrowhead indicates a translucent portion of a grain. (A) ZHE 733, (B) Nip, (C) N22, (D) Taggart, (E) Diamond, and (F) LaGrue.

**Figure 3:**
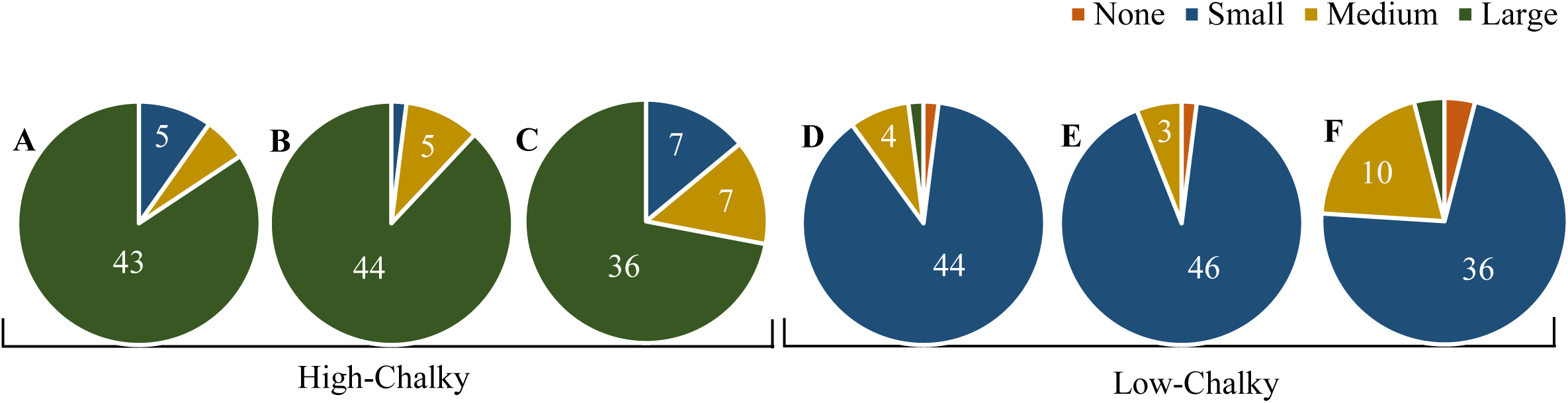
Distribution of chalk sizes. Chalk sizes were classified relative to the area of the grain. None (no chalk), small (less than 10%), medium (11-20%), and large (more than 20%). Scale adopted from Standard Evaluation System for Rice (SES), 2002. (A) ZHE 733, (B) Nipponbare, (C) Nagina 22, (D) Taggart, (E) Diamond, and (F) LaGrue. Numbers are frequency counts from 50 grains observed.

### Expression patterns of *AGPL4*, *GBSSI* and *SSIIA*

To understand how dynamics of starch biosynthesis controls grain chalkiness, gene expression analysis was performed in the spikelets of each line at six reproductive stages (BP, EF, 5DAF, 10DAF, 15DAF and 20DAF). Previous studies showed a direct correlation of mRNA abundance and enzyme activities of *AGPase*, *GBSS* and *SS* (Devi et al. 2010; Ponnala et al. 2014); therefore, gene expression directly correlates with the protein abundance and informs the dynamics of the starch biosynthesis process. High-chalky lines (ZHE 733, Nipponbare, Nagina 22) showed a characteristic pattern consisting of peak *AGPL4* expression at BP or AP with *GBSSI* and *SSIIA* peaks at 10DAF or 20DAF (Figure 4A-C), showing a temporal gap of ~15 days. Notably, in all three lines, *GBSS1* upregulation preceded that of *SSIIA*. ADP-Glucose, the product of AGPase, is unstable by nature (Baratt et al. 2009; Baroja-Fernandez et al. 2012); therefore, formation of ADP-Glucose before endosperm development would likely result in its non-utilization and reversal to Glucose-1-Phosphate. Further, since *AGPL4* is expressed at relatively lower levels during grain filling stages (Figure 4A-C), only a limited pool of ADP-Glucose, the substrate for GBSS and SS, is presumably available to synthesize starch components, amylose and amylopectin. Concurring with these observations, a previous study showed that uncoordinated expression of *AGPase* subunit genes (*AGPL3, AGPL4* and *AGPLS2*), subsequent to the peak expression of GBSS and SS, was associated with the inferior grains (Sun et al. 2015).

**Figure 4:**
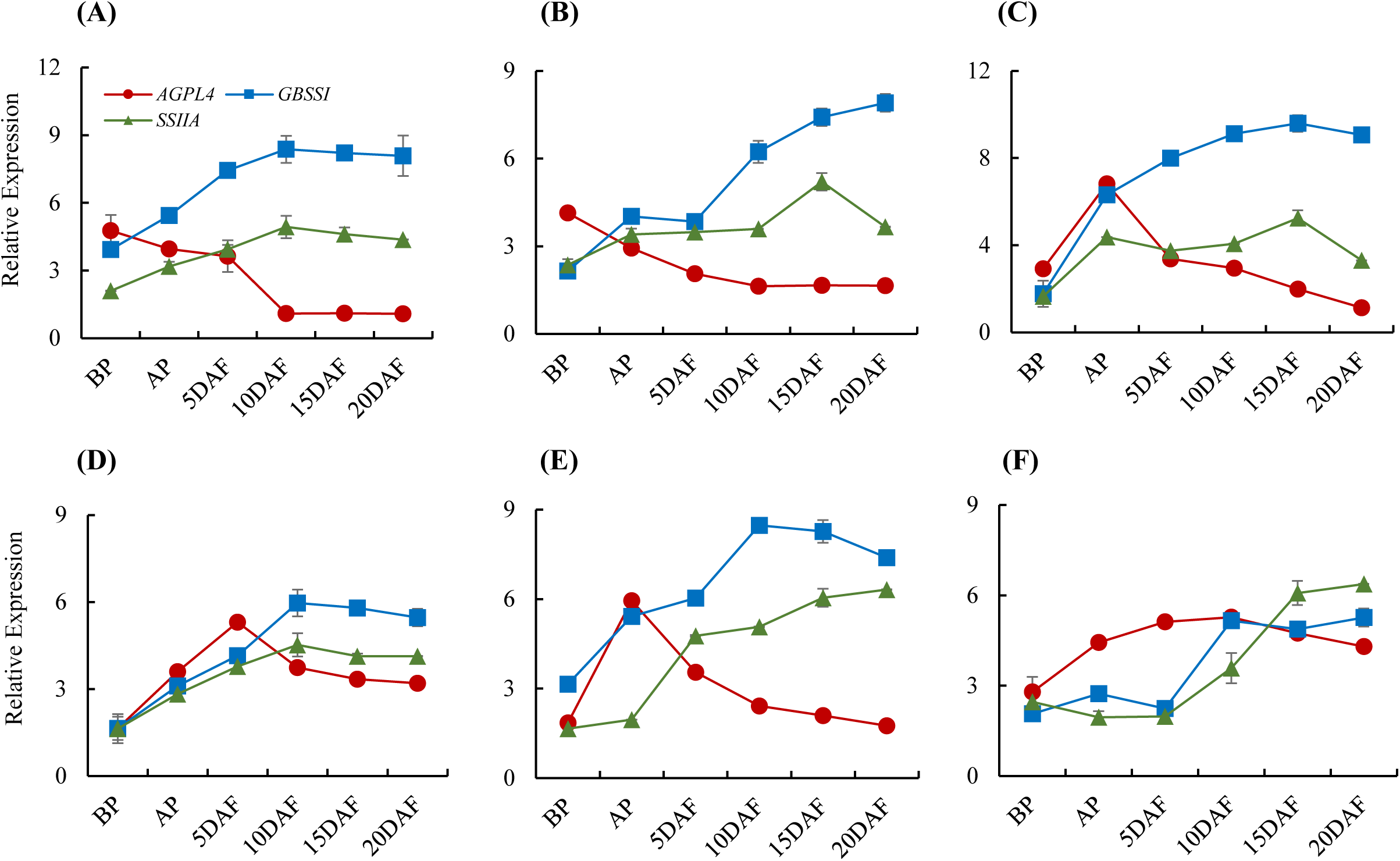
Gene expression patterns. Expression patterns of *AGPL4, GBSSI and SSIIA* during reproductive phases in rice with reference to the *7UBIQ* expression. (A) ZHE 733, (B) Nipponbare, (C) N22, (D) Taggart, (E) Diamond, and (F) LaGrue.

The three cultivars, on the other hand, showed a more coordinated pattern of expression characterized by the peak-expression of *AGPL4* followed quickly by the upregulation of *GBSSI* and *SSIIA* (Figure 4D-F). In Taggart, *AGLP4* peak occurred at 5DAF, followed immediately by the peaks of *GBSSI* and *SSIIA* at 10DAF (Figure 4D). Therefore, a sufficient pool of ADP-Glucose is presumably available for GBSS and SS to efficiently synthesize amylose and amylopectin, respectively, in the early phases of grain filling. This coordinated process leads to proper loading of starch in the form of edged granules, eventually forming non-chalky translucent grains. Another cultivar, Diamond, shows somewhat shifted expression peaks with *AGPL4* spike at AP that is quickly followed by the increase in both *GBSSI* and *SSIIA* activities at 5DAF and subsequent stages (Figure 4E). Thus, in Diamond, AGPase accumulates just before fertilization, and efficient starch synthesis commences soon after at 5DAF, possibly utilizing the pool of ADP-Glucose produced at the AP stage. Finally, LaGrue shows extended upregulation of *AGPL4* until 10DAF with gradual decline in the subsequent stages. This pattern either coincides or is immediately followed by *GBSSI* and *SSIIA* expression (Figure 4F), presumably allowing efficient conversion of ADP-Glucose to amylose and amylopectin. Finally, *GBSSI* expression is relatively higher than *SSIIA* at their peaks in all genotypes except LaGrue (Figure 4F) that showed higher *SSIIA* than *GBSSI* at 20DAF. This suggests active synthesis of amylopectin at this stage in LaGrue. However, as shown in Figure 1 and elucidated by others, amylopectin can be converted to amylose by starch debranching enzyme (*SDBE*) at later stages (Cheng et al. 2005).

In summary, the coordination between *AGPL4* and the two starch synthase genes, *GBSSI* and *SSIIA*, was evident in the three cultivars (low-chalk lines), while a gap of ~15 days between the peak expression of *AGPL4* and the starch synthase genes was observed in the high-chalky lines. This temporal gap may result in futile ADP-Glucose reaction, and a limited pool during the critical stages of grain filling.

### Granular morphology

SEM images showed differences in the granular morphology in high-chalky and low-chalky lines (Figure 5). Cross-section of the grains of high-chalky lines revealed simple granules of spherical (Figure 5A) or polyhedral shape (Figure 5B-C). Simple granules are mostly small in size, which points to early termination of amylose and amylopectin elongation. This hypothesis is supported by the expression analysis (Figure 4A) that suggested insufficiency of ADP-Glucose for amylose and amylopectin synthesis at the early grain filling stages. Simple granules allow more air spaces, which further elucidates the chalkiness of these genotypes. The high-chalky lines, Nipponbare and Nagina 22, show polyhedral granules of simple and compound types (Figure 5B-C). However, the sizes of the compound granules are not uniform. This heterogeneity of granules results in loose and irregular packing that explains the chalkiness in these genotypes. Protein bodies, as described by Kasem et al. (2011), are also observed in between and at the surface of the amyloplasts of ZHE 733 and Nipponbare, respectively (Figure 5A-B).

**Figure 5:**
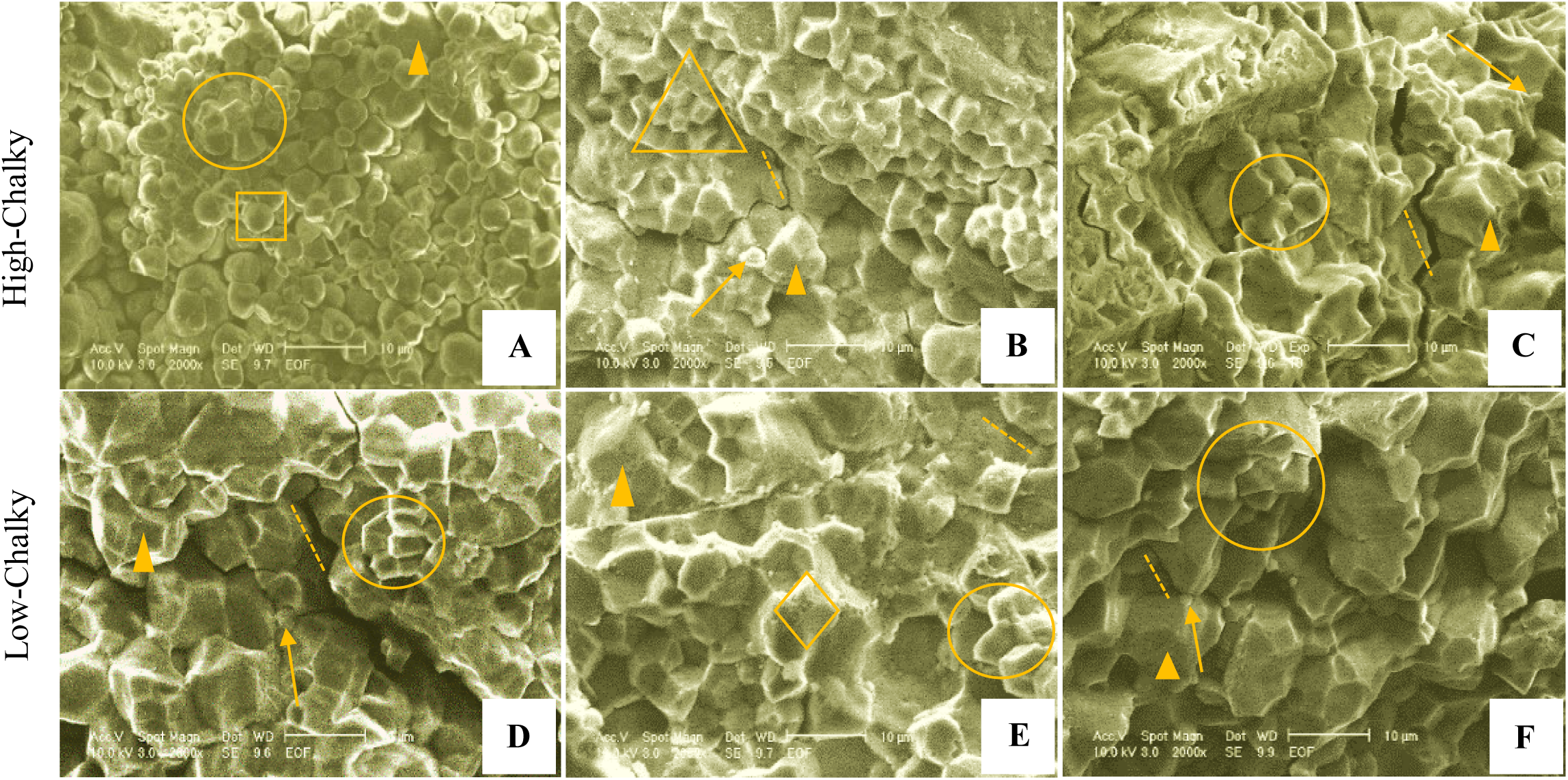
Granular morphology. Morphology of the grain transverse cross-section under scanning electron microscopy (SEM) with a scale bar of 10μm. (A) ZHE 733, (B) Nipponbare, (C) Nagina 22, (D) Taggart, (E) Diamond, and (F) LaGrue. Arrowhead indicates the surface of amyloplast. Rings show polyhedral granules inside split amylopast. Square indicates simple spherical granule. Triangle indicates simple polyhedral granules. Arrow points to protein bodies. Dashed line indicates the cracks from cross-sectioning the samples. Diamonds show micropores on the surface of amyloplast, which are the evidence of degradation activities.

Grains from low-chalky lines (cultivars) showed compact granular structure (Figure 5D-F). These granules appear compound type, suggesting their origin from efficient chain elongation and biosynthesis of starch (Toyosawa et al. 2016). Compound granules are large and can be an aggregate of smaller granules packed very tightly. This kind of granules is produced when substrates and enzymes are sufficiently available at the critical stages of grain filling. These conditions appear to be fulfilled in the low-chalky cultivars, as indicated by the coordinated expression pattern of starch synthesis genes (Figure 4D-F). Moreover, the granules are homogeneous affording tight packing and preventing air spaces in the grains. Further, smaller protein bodies were observed in the cultivars. However, micropores were observed in the amyloplastic surface of Diamond, indicating possible degradation by α-amylases (Figure 5E-F). Similar granular structures were observed in previous studies on the chalky grains (Lin et al. 2016; Kaneko et al. 2016; and Mitsui et al. 2016).

### Biochemical analysis

Higher amylose content was observed in the high-chalky lines as compared to the low-chalky lines (Figure 6). Within the high-chalky group, the indica rice ZHE 733, that shows simple and spherical granules (Figure 5A), contained significantly higher amylose fraction (Figure 6A). Although, physicochemical properties of the indica and japonica rice may differ, high amylose cultivars have generally been reported to show simple granular morphology under SEM (Zhang et al. 2017). A possible mechanism underlying this phenotype is the premature termination of α-glucan chain elongation due to insufficiency of ADP-Glucose at the grain filling stages as suggested by the *AGPL4* expression pattern (Figure 4A). As a result, shorter chains are generated and branching (1 **→** 6) in the amylopectin is reduced, leading to simple granules. This analogy is reinforced by observations that high amylose rice grains contain shorter α-glucan chain length (Park et al., 2007). However, SDBE mediated debranching of amylopectin can also lead to higher amylose (Cheng et al. 2005; Figure 1).

**Figure 6:**
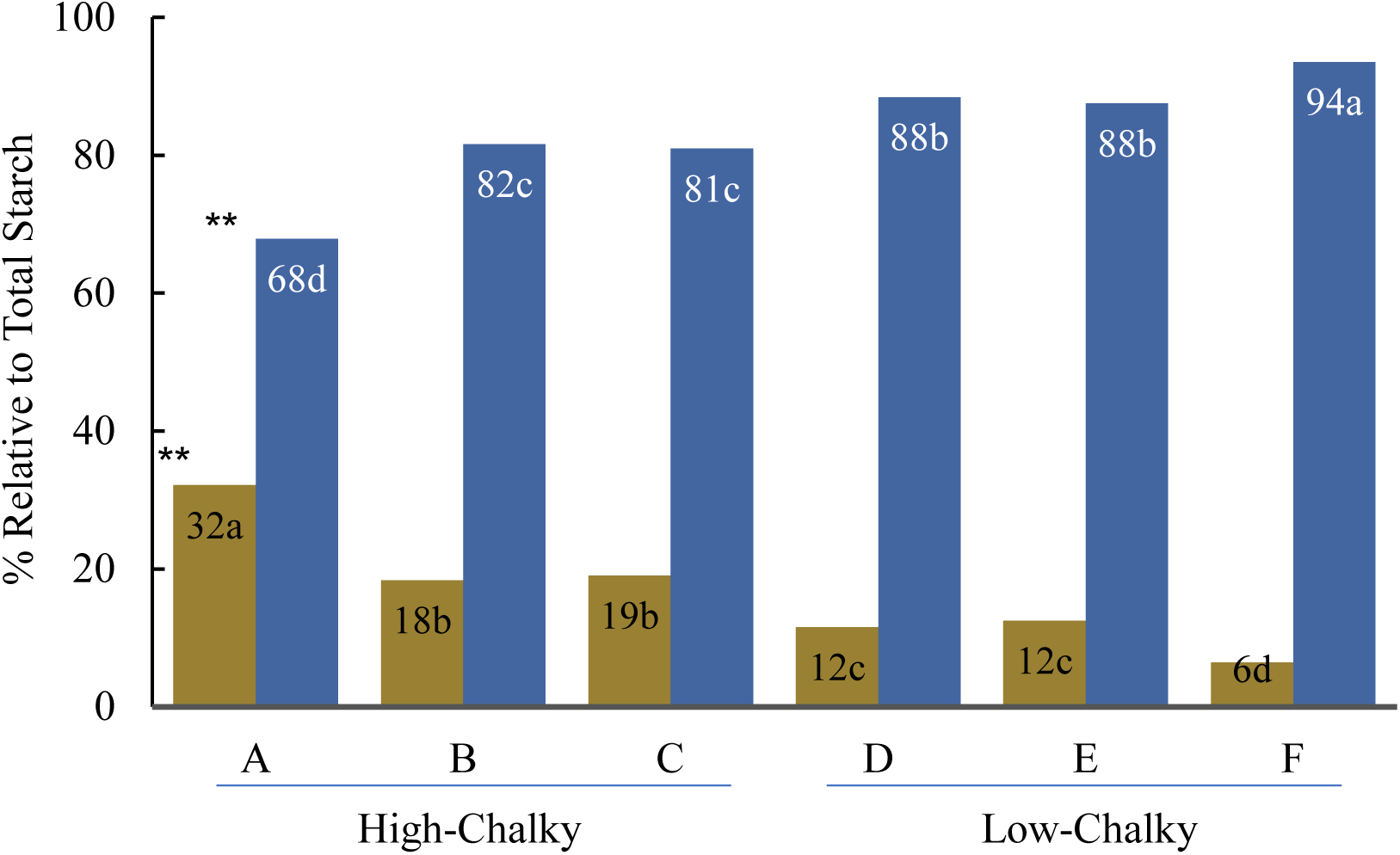
Amylose and amylopectin content (%) relative to the total starch. Top=amylose and botton=amylopectin. Asterisks indicate significant differences among the genotypes (**at p=0.01). Lower case letters in each bar are the ranks from mean comparison. (A) ZHE 733, (B) Nipponbare, (C) N22, (D) Taggart, (E) Diamond, and (F) LaGrue

Low-chalky cultivars were found to contain significantly higher amylopectin content (Figure 6D-E), which corroborates with a previous study that found the association of higher amylopectin with lower chalkiness (Lin et al. 2016). Higher amylopectin can be attributed to an efficient starch biosynthetic process and longer α-glucan chain lengths. This efficiency arguably relies on the coordinated expression of *AGLP4*, *GBSSI* and *SSIIA* as suggested by the gene expression analysis on the low-chalky cultivars (Figure 2D-F).

Regardless of the chalkiness, amylopectin is higher than amylose in all lines despite lower expression of *SSIIA* relative to *GBSSI* (Figure 4). This suggests that other isoforms of SS actively participate in the catalysis of the same biochemical pathway to synthesize amylopectin, especially *SSI* that is upregulated during 28-42DAF (Figure 1). Accordingly, *SSI* mutants show decrease in amylopectin content (Abe et al. 2014). However, starch branching enzymes (SBE) also contributes to the pool of amylopectin, and mutagenesis of *SBEIIB* by CRISPR/Cas9 decreased amylopectin in rice grains (Sun et al. 2017).

Higher protein content has also been linked to higher chalkiness (Yamakawa & Hakata, 2010). Nipponbare and Nagina 22, the two high-chalky lines, were found to have highest protein concentrations (Figure 7), suggesting that higher number of protein bodies in these genotypes could contribute to grain chalkiness. ZHE 733, on the other hand, showed a similar level of protein content as the three cultivars, suggesting proteins bodies do not play a major role in the chalkiness of ZHE 733.

**Figure 7:**
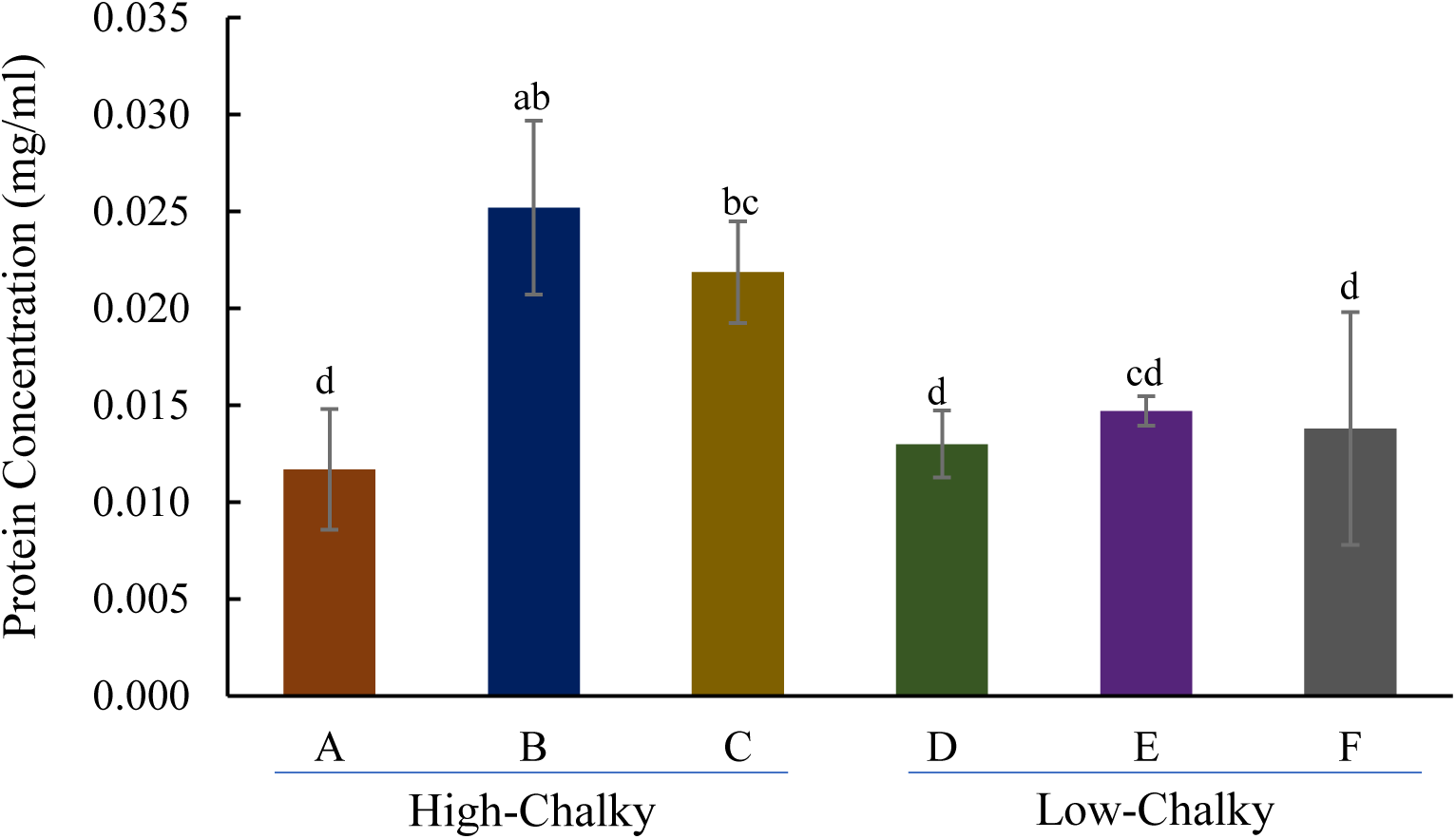
Protein content in the grains of different genotypes. Significant at α=0.05, bars with different letter designations are not comparable under Tukey’s mean comparison. (A) ZHE 733, (B) Nip, (C) N22, (D) Taggart, (E) Diamond, and (F) LaGrue.

### Effect of heat stress on the expression patterns

Heat stress has a major effect on grain chalkiness. Under high nighttime temperature (HNT), there is increased formation of chalk accompanied with the changes in the expression of starch biosynthesis genes (Lanning et al. 2010, Nevame et al. 2018; Dhatt et al. 2019). To determine whether alteration in the expression patterns is a part of the mechanism associated with the HNT-induced chalkiness, three genotypes, ZHE 733, Diamond, and LaGrue were analyzed under HNT conditions. Of these three, ZHE 733 is chalky in the normal condition without heat treatment (Esguerra et al. 2019).

Upon HNT treatment, all three genotypes showed altered expression of *AGPL4*, *GBSSI* and *SSIIA*. In the cultivar Diamond, the most striking alteration occurred in *AGPL4* expression pattern, which was upregulated at BP but downregulated in subsequent stages (Figure 8A). This concurs the expression pattern of other AGPase subunit genes in plants under HNT stress. Dhatt et al. (2019) reported that *AGPS2* and *AGPL2* are downregulated during 7-10DAF and upregulated at 4DAF under HNT condition. In LaGrue, *AGPL4* spiked at an earlier stage (AP) by HNT with a rapid decline in the subsequent stages. Whereas in the normal condition, it is gradually increased during early grain filling stages (Figure 8B). Additionally, *GBSSI* and *SSIIA* in LaGrue were markedly elevated by HNT, which may lead to hyperactivity of these enzymes and depletion of ADP-Glucose in subsequent stages. Finally, ZHE 733, which is equally chalky in the normal and HNT conditions (Esguerra et al. 2019), showed a similar expression pattern of the three genes. The relative expression of *AGPL4* is elevated and that of *GBSSI* and *SSIIA* is suppressed by HNT for ZHE 733 (Figure 8C).

**Figure 8:**
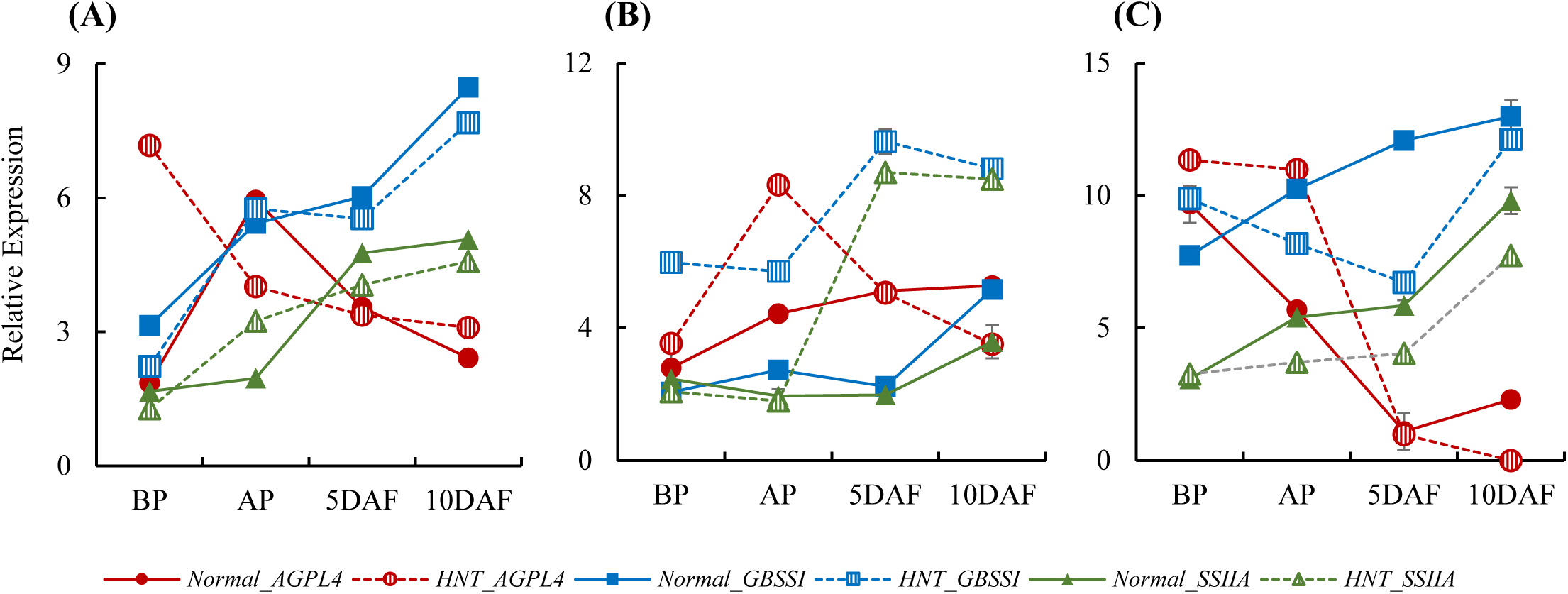
Effect of heat stress on gene expression. Relative expression of AGPL4, GBSS1, and SSIIA in normal and high nighttime temperature (HNT) conditions in (A) Diamond, (B) LaGrue and (C) ZHE 733 with reference to the *7UBIQ* expression. Solid lines indicate normal condition and dashed lines indicate HNT

Next, physical, granular, and chemical properties of grains in Diamond and LaGrue were analyzed. The chalk distribution and grain morphology in the two cultivars completely changed upon HNT treatment. In the normal condition, these cultivars produced low-chalky grains; however, under HNT, both cultivars developed large chalky areas (Figure 9A-B). Interestingly, the granular morphology under HNT was different in the two cultivars. The HNT-Diamond developed simple, spherical granules (Figure 9C); whereas HNT-LaGrue retained compound polyhedral granules. However, HNT-LaGrue developed many shreds and micropores (Figure 9D). Notably, the granular morphology of HNT-Diamond resembled that of ZHE 733 (Figure 5A, Figure 9C). Interestingly, the expression patterns of starch synthesis genes in HNT-Diamond was also similar to that of ZHE 733 (Figure 4A, Figure 8A). The formation of compound polyhedral granules in HNT-LaGrue is consistent with the upregulation of *AGPL4*, *GBSSI*, and *SSIIA* (Figure 8B). However, abundance of shreds and micropores in HNT-LaGrue points to amylase attack as part of the mechanism.

**Figure 9:**
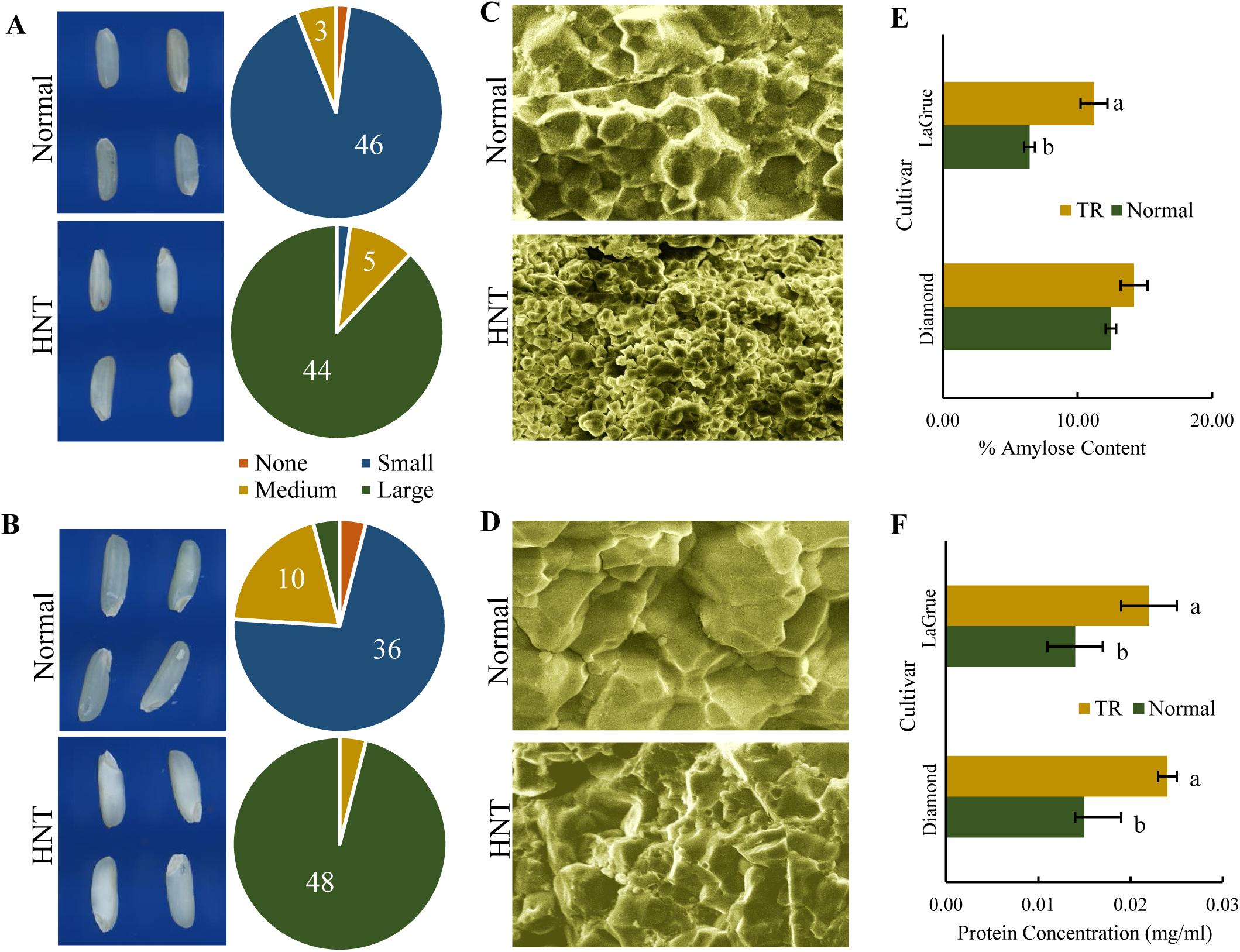
Physicochemical analysis of grains under normal and high nighttime temperature (HNT) conditions. (A-B) Grain images captured using WINSEEDLE and distribution of chalk sizes in Diamond (A) and LaGrue (B). (C-D) Granular morphology under scanning electron microscope (SEM) of Diamond (C) and LaGrue (D). (E) Amylose and amylopectin content in normal and HNT condition. (F) Protein concentration in normal and HNT condition.

Finally, amylose and protein contents were determined under normal and HNT conditions. In Diamond, no significant difference in the amylose content was observed between the two conditions, while in LaGrue an increase in the amylose content was observed in the HNT grains (Figure 9E). Previous studies have reported both increase or decrease in amylose content in response to HNT (Ahmed et al. 2014; Cheabu et al. 2018). However, the protein content in both cultivars increased significantly under HNT (Figure 9F). Increased protein content has been associated with higher number of protein vesicles that contribute to the grain chalk (Kaneko et al. 2016).

## CONCLUSIONS

In conclusion, coordinated expression of starch biosynthesis genes is important in developing compound polyhedral granules associated with the non-chalky translucent rice grains. Perturbation of this process by HNT leads to altered granular structure conferring grain chalkiness. Other factors such as number of protein bodies and amylase activities also play a significant role in grain chalkiness, apparently in a genotype-dependent manner. Nevertheless, in the early phases of grain development, efficient synthesis of starch through the coordinated activities of AGPase, GBSS and SS is arguably the most important mechanism controlling granular morphology in a genotype-dependent manner. In the coordinated expression pattern, AGPase is upregulated early in the reproductive phase, closer to the grain filling stages, to produce abundant pool of ADP-Glucose, which is quickly utilized by GBSS and SS to produce amylose and amylopectin in amyloplasts. This streamlined process leads to uniform polyhedral granules that pack tightly and produce non-chalky grains (Figure 10A). However, when AGPase is upregulated too early in the reproductive phase, the resulting ADP-Glucose reverts back to Glucose-1-P. As a result, only a limited pool of ADP-Glucose is presumably available for starch synthesis during the early grain filling stages. This uncoordinated process could lead to early termination of chain elongation, producing smaller granules of heterogeneous shapes (Figure 10B). These simpler spherical or heterogeneous granules pack more loosely and accommodate air spaces observed as chalk in the mature grains.

**Figure 10:**
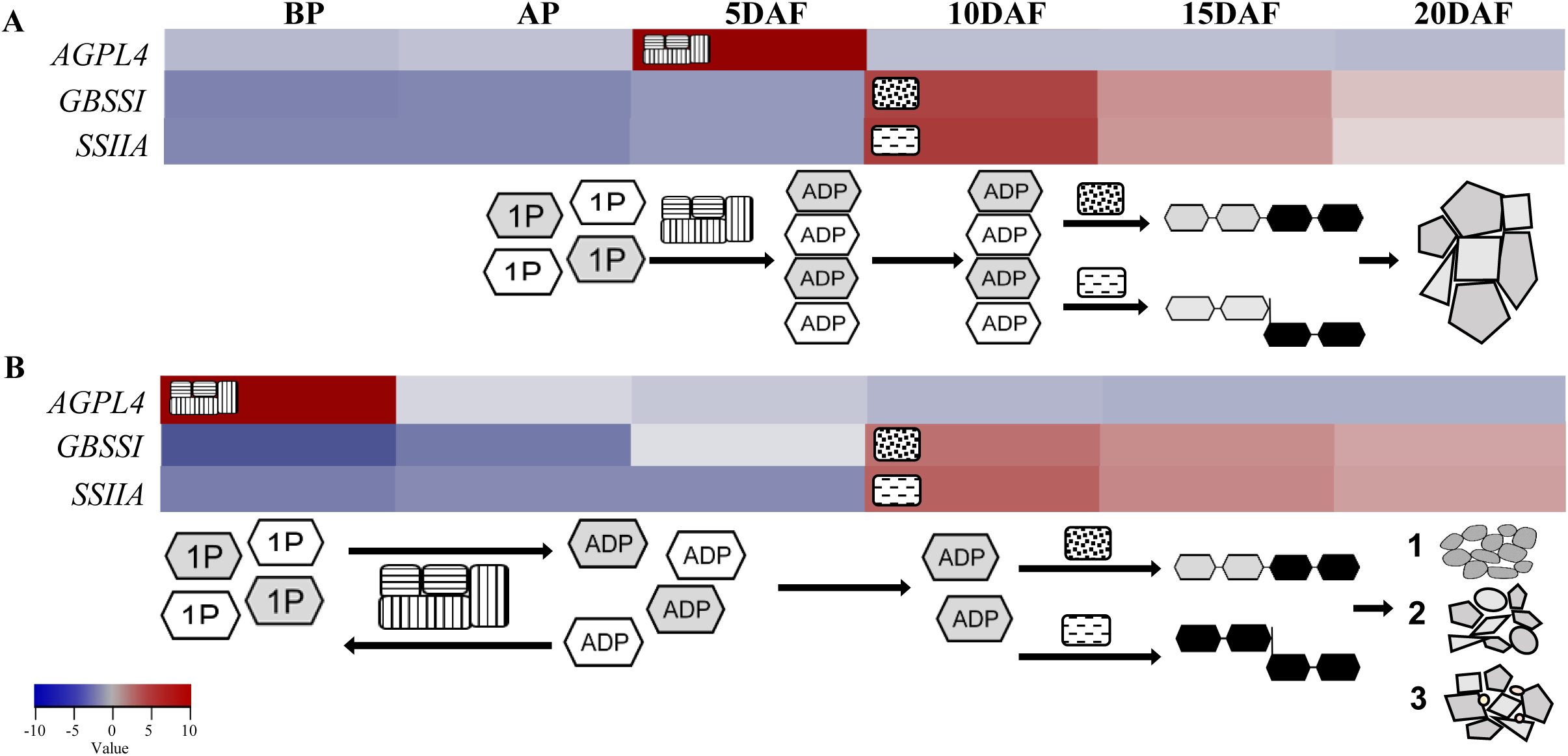
Proposed mechanism of grain chalkiness. (A) Coordinated, and (B) Uncoordinated expression of *AGPL4*, *GBSSI* and *SSIIA* through early reproductive phases. (A) AGPL4, the regulatory unit of tetrameric AGPase, and monomeric GBSS1 and SSIIA are upregulated in quick succession through early grain filling stages. As a result, the conversion of Glucose-1-Phosphate (1P) to ADP-Glucose (ADP) occurs in a timely manner to efficiently synthesize amylose and amylopectin. This coordinated expression leads to the formation of large polyhedral granules packed tightly in the grains. (B)A temporal gap between the upregulation of AGLP4 and that of GBSS1 and SSIIA leads to non-utilization of ADP, resulting in the reversal of ADP to 1P. When GBSS1 and SSIIA are upregulated in subsequent stages, the limited pool of ADP may lead to early termination of chain elongation. This uncoordinated process leads to smaller granules of spherical or polyhedral shapes accommodating airspaces and protein bodies that appear as grain chalk. The heatmap represents developmental expression pattern of the high-chalky (Figure 2A) and low-chalky line (Figure 2D) through reproductive stages BP, AP, 5DAF, 10DAF, 15 DAF, and 20DAF. Linear chain of hexagons indicate amylose (α1 **→** 4 glycosidase linkage) and branched chain indicates amylopectin (α1 **→** 4 & 1 **→** 6 glycosidase linkage).

## Notes

### Competing Interest Statement

The authors have declared no competing interest.

